# Intermittent water stress favors microbial generalists that better help wheat under drought

**DOI:** 10.1101/2023.11.16.567418

**Authors:** Ruth Lydia Schmidt, Hamed Azarbad, Luke Bainard, Julien Tremblay, Etienne Yergeau

## Abstract

Microorganisms can improve plant resistance to drought through various mechanisms such as the production of plant hormones, osmolytes, antioxidants, and exopolysaccharides. It is, however, unclear how previous exposure to water stress affects the functional capacity of the soil microbial community to help plants resist drought. We compared two soils that had either a continuous or intermittent water stress history for almost forty years. We grew wheat in these soils and subjected it to a water stress, after which we collected the rhizosphere soil and shotgun sequenced its metagenome. Wheat growing in the soil with an intermittent water stress history maintained a higher fresh biomass when subjected to water stress. Genes related to resistance to drought were more abundant in the metagenome and more prevalent, diversified, and redundant in the metagenome assembled genomes of the soil with an intermittent water stress history as compared to the soil with a continuous water stress history. We suggest that an intermittent water stress history selects for generalists that are adapted to both low and replete water contents, and that these generalists harbor a larger repertoire of genes beneficial for life under water stress.

## Introduction

Plant-and soil-associated microorganisms play a pivotal role in mitigating yield losses due to drought. These microorganisms have evolved diverse mechanisms to resist or avoid drought, which can also be beneficial to plants [1]. Unlike avoidance mechanisms like dormancy or sporulation, microbial resistance mechanisms allow microorganisms to remain active and support plants during drought. Key resistance mechanisms include osmolyte production to retain water within cells [2], extracellular polymeric substances (EPS) production to enhance soil water retention [3], and production of various enzymes to detoxify reactive oxygen species (ROS) generated under stress [4]. These mechanisms not only help microorganisms but also have positive effects on plants during drought. For example, microbially produced osmolytes can be transferred to plant tissues [5, 6], and microbially produced EPS near plant roots can enhance water-holding capacity [7]. Microorganisms also modulate the plant’s hormonal response to stress. The microbial 1-aminocyclopropane-1-carboxylate (ACC) deaminase degrades the precursor of the stress hormone ethylene [8]. Furthermore, the production of auxins and cytokinins by microorganisms can promote plant growth and resistance to stress [9, 10].

Although interfering with the regular plant stress response might offer short-term benefits, it could have long-term consequences, especially since larger plants are more susceptible to drought [11]. The overall beneficial impact of microbial activities on crops during drought depends on factors such as the prevalence, abundance, diversity, and expression of traits, which, in turn, are influenced by various biotic and abiotic factors, including water availability.

Microbial communities respond to actual water availability in their environment, either through resistance or avoidance mechanisms. This selective pressure, when sustained or repeated over time, can lead to lasting shifts in microbial community composition and activities. The frequency of the stress event is also critical, as intermittent stress selects for generalists adapted to both stressful and normal conditions, whereas continuous stress favors microorganisms specialized for life under stressful conditions [12, 13]. A case in point, the microbial communities in two adjacent dryland wheat field soils subjected to intermittent or continuous water stress over nearly 40 years not only differed in composition but also responded differently to water stress [14, 15]. When wheat grew in soil with a continuous history of water stress, root biomass decreased more sharply when subjected to a subsequent water stress compared to roots in soil with an intermittent water stress history [16]. Additionally, microbial communities extracted from soil with an intermittent water stress history reduced catalase activity in the leaves (indicative of lower stress levels) when inoculated into wheat growing in different soil and subjected to water stress [17]. Since these studies relied on amplicon sequencing, it remains unclear how the differences observed in microbial communities of the soils historically subjected to different frequencies of stressful events translate to variations in microbial traits and their impact on plant stress resistance.

In this study, we used shotgun-sequencing to analyze the rhizosphere metagenome of wheat plants from the soils of Azarbad et al. [15, 16], which had continuous or intermittent water stress histories. Our hypothesize was that intermittent water stress, due to its variable selection pressure, would favor a greater prevalence and diversity of generalist soil microbes with functional traits related to surviving at low water availability, contrasting with the constant water stress history. This increased diversity and prevalence of generalist microorganisms is expected to enhance plants’ resistance to water stress events.

## Material and methods

### Experimental design

We utilized samples from the pot study conducted by Azarbad et al. [15, 16]. The soil utilized in the pots was sourced from two adjacent fields that had been subjected to distinct irrigation since 1981. Both fields followed wheat-fallow two-year rotations, but one field was irrigated during the wheat phase of the rotation, while the other remained unirrigated. Since the fields are in the semi-arid region of Saskatchewan, near the Swift Current Research and Development Centre (Agriculture and Agri-Food Canada), this difference in irrigation resulted in continuous and intermittent water stress conditions. For the current metagenomic study, we used rhizosphere soil derived from one drought-sensitive wheat cultivar (*Triticum aestivum* cv. AC Nass), which was cultivated in one of the two soil types at 5% or 50% soil water holding capacity (SWHC). These treatments were chosen among the entire experimental design based on previous studies that showed large contrasts in microbial communities [14–16]. This resulted in a total of four treatments with five replicates each, yielding a total of 20 samples. Each pot contained 1 kg of soil and was seeded with eight seeds. For the first 4 weeks, soil water content was kept at 50% SWHC for all pots, after which they were either kept at 50% SWHC or brought to 5% SWHC for another 4 weeks. At the end of the experiment, plants were vigorously shaken and the rhizosphere soil that remained attached to the roots was then harvested, flash-frozen in liquid nitrogen and kept at −80°C until DNA extraction. Additional details regarding the soil composition and the pot experiment can be found in Azarbad et al. [14–16].

### DNA extraction and sequencing

The DNA was extracted using a DNeasy PowerSoil kit (Qiagen) and sent for metagenomic sequencing at the Centre d’expertise et de services Génome Québec located in Montréal, Québec. The sequencing procedure performed using an Illumina HiSeq 4000 (PE150), yielded a total of 699,061,058 reads, with an average of 34,953,053 reads per sample, resulting in a total of 105 Gbp, or an average of 5.2 Gbp per sample. The raw data has been deposited under BioProject accession PRJNA1040208.

### Bioinformatics

The sequencing reads were processed using our established metagenomic pipeline (ShotgunMG v 1.3.2), as previously described [18, 19]. Quality-controlled reads from each sample were co-assembled, after which genes were identified and annotated on each assembled contig. Subsequently, reads were mapped oton the contigs to derive abundance profiles which were used as input to generate Metagenome-Assembled Genomes (MAGs) (MetaBat v 2.12.1).

### Functional traits

For the functional trait analyses, we searched in our gene annotation table for entries related to the functions of interest using their KEGG orthology (KO) entries. For the ACC deaminase we used the only KO available for this function: K01505. For IAA production, we used KOs of enzymes that led directly to IAA in the tryptophan metabolism map (map00380): K01501, K01426, K21801, K11816, K11817, K00128. For osmolytes production we used the KO list present in the supplementary table 2 of McParland et al. [20]. For EPS biosynthesis, we used the 73 KOs associated with the KEGG pathway ko00543 (Exopolysaccharide biosynthesis). For cytokinin, we used the 10 KOs associated with the KEGG pathway ko00908 (Zeatin biosynthesis). For antioxidants, we searched for KOs with the terms “superoxide dismutase” (5 entries), “glutathione peroxidase” (4 entries), “catalase” (4 entries), or “cytochrome oxidase” (58 entries).

### Statistics

All R code used for data manipulation, statistical analyses, and figure generation can be found on our GitHub repository (https://github.com/le-labo-yergeau/MG_Growth_Room). The data employed in the R scripts have been deposited on the Zenodo platform (https://zenodo.org/doi/10.5281/zenodo.10140592).

## Results

### Plant biomass and leaf water content

In comparison to the high soil water content (50% soil water holding capacity) treatment (HWC), the low soil water content (5% soil water holding capacity) treatment (LWC) reduced plant fresh biomass by 83.8% (roots) to 84.6% (shoots), on average (P<0.001, Table 1, Fig. 1a). For shoots, this reduction in biomass can be attributed, at least in part, to a 59.7% reduction in leaf water content (P<0.001, Table 1). Soil water stress history (WSH) also had an impact on shoot biomass, with shoot biomass being reduced by 10.8% in the continuous WSH soil as compared to the intermittent WSH soil, which is most evident under LWC (P=0.028, Table 1, Fig. 1a). Additionally, the root biomass of HWC plants was 49.3% higher for plants growing in the soil with a continuous WSH compared to the plant growing in the soil with an intermittent WSH, but this was not significant under LWC (interaction term: P<0.001, Table 1, Fig. 1a).

**Figure 1.**
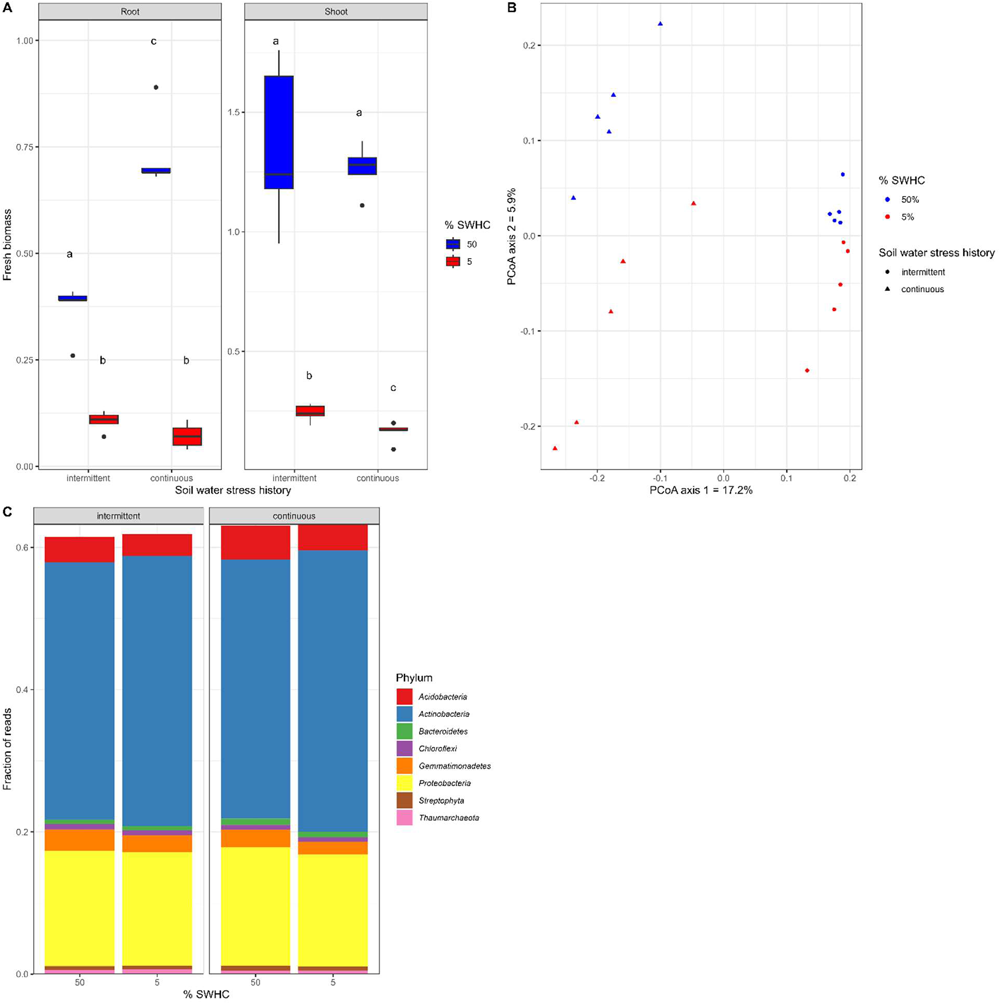
Plant and general microbial response. A) root and shoot fresh biomass, B) metagenomic community structure using a principal coordinate analysis of Bray-Curtis dissimilarities, and C) metagenomic community composition for rhizosphere samples taken from wheat growing in soil with an intermittent or continuous water stress history and subjected to low (5% soil water holding capacity) or high (50% soil water holding capacity) water availability. Different letters in A) indicate significant differences at P<0.05.

**Table 1.** Anova table for plant root and shoot fresh biomass, and leaf water content for wheat subjected to water stress and growing in two soils with contrasting soil water stress history.

|  | Roots |  | Shoots |  | Leaf water content |  |
| --- | --- | --- | --- | --- | --- | --- |
|  | F | P | F | P | F | P |
| Soil type | 1.13 | 0.31 | 6.22 | 0.028* | 2.02 | 0.18 |
| % SWHC | 229.12 | 3.5x10<sup>-9</sup>*** | 397.47 | 1.45x10<sup>-10</sup>*** | 153.17 | 3.43x10<sup>-8</sup>*** |
| Soil:SWHC | 21.72 | 0.00055*** | 3.98 | 0.069. | 0.420 | 0.53 |

\*\*\*:P<0.001; \*\*: 0.001<P<0.01; \*: 0.01<P<0.05; .: 0.05<P<0.10.

Neither WSH nor the interaction term affected the leaf water content (P>0.05, Table 1), suggesting that their effect on leaf fresh biomass was not due to changes in water content.

### Microbial community composition

The gene community structure showed that rhizosphere samples from wheat growing in the same soil were more similar to each other than to the other soil (PCoA of Bray-Curtis dissimilarity: Fig 1b and Permanova: Table 2, F=6.98, P=0.0001). The current soil water content also resulted in clustering of the wheat rhizosphere samples (Fig. 1b), but this was not significant in Permanova tests (F=1.45, P=0.16, Table 2). The community composition at the phylum level did not differ (Fig. 1c), suggesting that the two soils were taxonomically similar at that level. The annotated reads were mainly affiliated with the Actinobacteria and, to a lesser extent, to the Proteobacteria (Fig. 1c).

**Table 2.** Permanova results for the composition of functional genes in the rhizosphere of wheat subjected to water stress and growing in two soils with contrasting soil water stress history.

|  |  | Soil - historical | %SWHC - current | %SWHC:SWSH |
| --- | --- | --- | --- | --- |
| All genes | F | 6.98 | 1.45 | 1.14 |
|  | P | 0.0001*** | 0.16 | 0.26 |
| IAA | F | 7.17 | 1.54 | 1.17 |
|  | P | 0.0001*** | 0.13 | 0.26 |
| ACC | F | 8.11 | 1.57 | 1.30 |
|  | P | 0.0001*** | 0.14 | 0.22 |
| Osmolytes | F | 7.76 | 1.44 | 1.14 |
|  | P | 0.0001*** | 0.17 | 0.28 |
| EPS | F | 7.20 | 1.41 | 1.11 |
|  | P | 0.0001*** | 0.17 | 0.28 |
| Cytokinines | F | 7.05 | 1.44 | 1.06 |
|  | P | 0.0002*** | 0.16 | 0.33 |
| ROS | F | 7.76 | 1.44 | 1.14 |
|  | P | 0.0001*** | 0.17 | 0.28 |

\*\*\*: $P < 0.001$ ; \*\*: $0.001 < P < 0.01$ ; \*: $0.01 < P < 0.05$ ; $\therefore 0.05 < P < 0.10$ .

### Drought-related traits

We compared the total abundance and the composition of six genes/pathways that could be involved in microbial beneficial services to the plants under water stress: indoleacetic acid (IAA) synthesis, ACC deaminase synthesis, cytokinin metabolism, extracellular polymeric substances (EPS) synthesis, osmolyte production and antioxidant synthesis. Like the patterns observed for all gene, the gene composition of the subgroups was only influenced by the WSH (P<0.0001 for all, Table 2). However, when looking at the summed abundance of the genes in a subgroup, different patterns emerged (Fig. 2, Table 3). For instance, the relative abundance of IAA and EPS related genes and of the ACC deaminase were influence by both the WSH and by the current soil water content; the relative abundance of osmolyte-related genes was only influenced by the WSH; the relative abundance of antioxidants was only influenced by actual soil water content; and the relative abundance of cytokinins was not influenced by any of the experimental factors (Table 3). Even if there were similitude in the factors affecting these group of genes, the patterns were not the same. The ACC-deaminase gene was 7% more abundant in the rhizosphere of wheat growing in soil with a continuous WSH, and it was also 5% more abundant in the LWC rhizospheres (Fig. 2a). IAA-related genes were 3% more abundant in the soil with an intermittent WSH as compared to the soil with a continuous WSH (Fig. 2b). Similarly, IAA genes were 1.6% more abundant in the LWC treatment as compared to the HWC treatment (Fig. 2b). Osmolyte genes were 2% more abundant in the rhizosphere of plants growing in the soil with an intermittent WSH (Fig. 2c). EPS production genes were 1.7% more abundant in the soil with a continuous WSH and 1.6% more abundant in the HWC pots (Fig. 2d). There was no significant difference for the cytokinins (Fig. 2e). Antioxidant-related genes were 1.3% more abundant in the soils with a continuous WSH and 2.7% more abundant in HWC soils (Fig. 2f).

**Figure 2.**
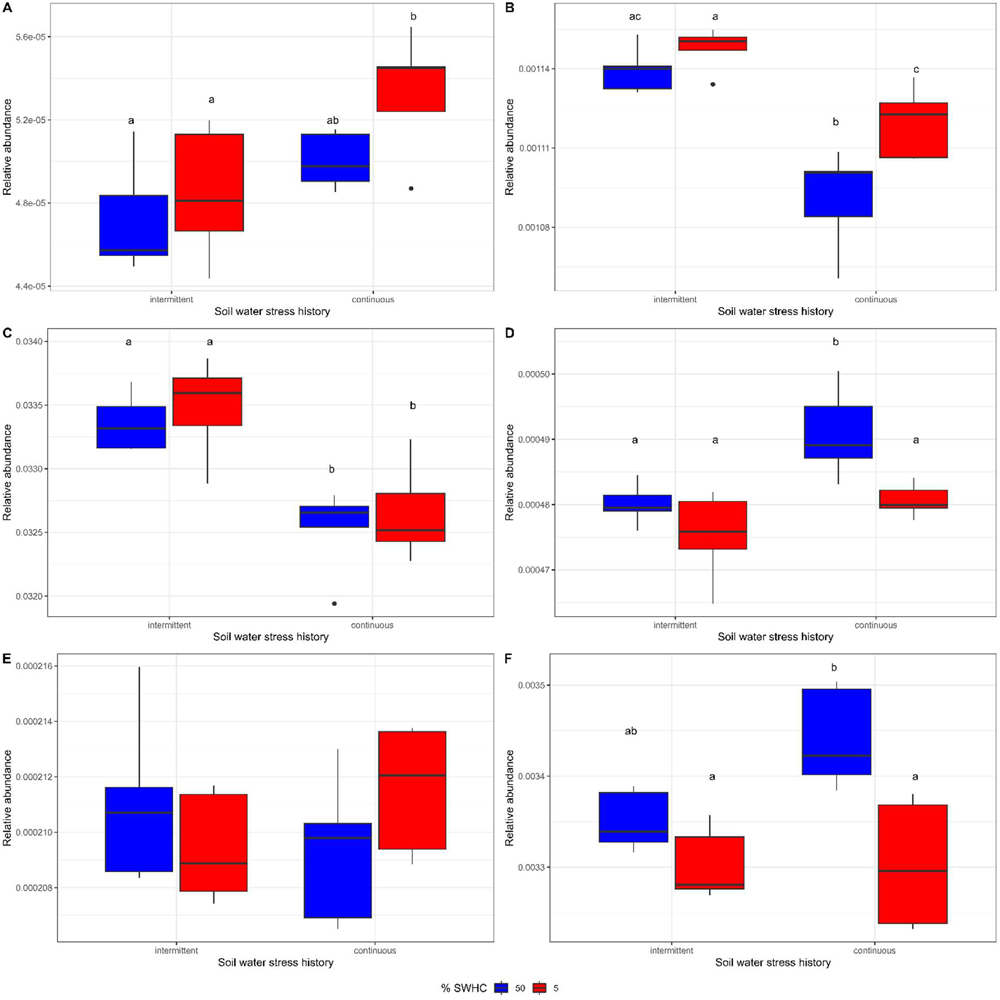
Relative abundance of genes encoding for drought-related functional traits. A) ACC deaminase, B) indole acetic acid, C) osmolytes, D) extracellular polymeric substances, E) cytokinins, F) antioxydants relative abundance for rhizosphere samples taken from wheat growing in soil with an intermittent or continuous water stress history and subjected to low (5% soil water holding capacity) or high (50% soil water holding capacity) water availability. Different letters indicate significant differences at P<0.05.

**Table 3.** Anova results for the relative amount of functional genes in the rhizosphere of wheat subjected to water stress and growing in two soils with contrasting soil water stress history.

|  |  | Soil - historical | %SWHC - current | %SWHC:SWSH |
| --- | --- | --- | --- | --- |
| IAA | F | 49.07 | 11.44 | 3.58 |
|  | P | 1.42x10<sup>-5***</sup> | 0.0055** | 0.083. |
| ACC | F | 14.06 | 5.00 | 0.95 |
|  | P | 0.0028** | 0.045* | 0.35 |
| Osmolytes | F | 35.71 | 0.76 | 0.001 |
|  | P | 6.45x10<sup>-5***</sup> | 0.40 | 0.98 |
| EPS | F | 19.66 | 17.02 | 2.22 |
|  | P | 0.00082*** | 0.0014** | 0.16 |
| Cytokinins | F | 0.025 | 0.084 | 3.02 |
|  | P | 0.88 | 0.78 | 0.11 |
| ROS | F | 3.48 | 14.71 | 3.54 |
|  | P | 0.087. | 0.0024*** | 0.084. |

\*\*\*: P<0.001; \*\*: 0.001<P<0.01; \*: 0.01<P<0.05; .: 0.05<P<0.10.

### Metagenome assembled genomes (MAGs)

Among the 714 MAGs assembled – containing approximately 30% of all reads – 434 were affected by the soil water stress history, 45 by the actual soil water content and 2 by the interaction term (Bonferroni corrected P<0.05/714). Among the 435 MAGs affected by soil history, 171 were enriched in the soil with intermittent WSH, and 264 were enriched in the soil with continuous WSH. As compared to the unaffected MAGs, the Actinobacteria were overrepresented in the MAGs enriched in the soil with a continuous WSH, whereas the Proteobacteria were underrepresented. Most functional traits were more prevalent in the MAGs that were affected by the soil WSH, as compared to the unaffected MAGs (Fig. 3a). Except for the ACC deaminase gene, the prevalence of functional traits was higher in the MAGs that were enriched in the soil with an intermittent WSH (Fig. 3a). A higher number of genes related to the different traits was also found in the MAGs that were affected by soil WSH (Fig. 3c). A higher percentage of the intermittent WSH soil MAGs encoded for more than 3 functional traits as compared to the unaffected or the continuous WSH soil MAGs (Fig. 3e). This means that intermittent soil WSH selected for bacteria not only harboring genes encoding functional traits related to life under water stress, but also for bacteria having several copies of these genes and that combined genes linked to several different traits.

**Figure 3.**
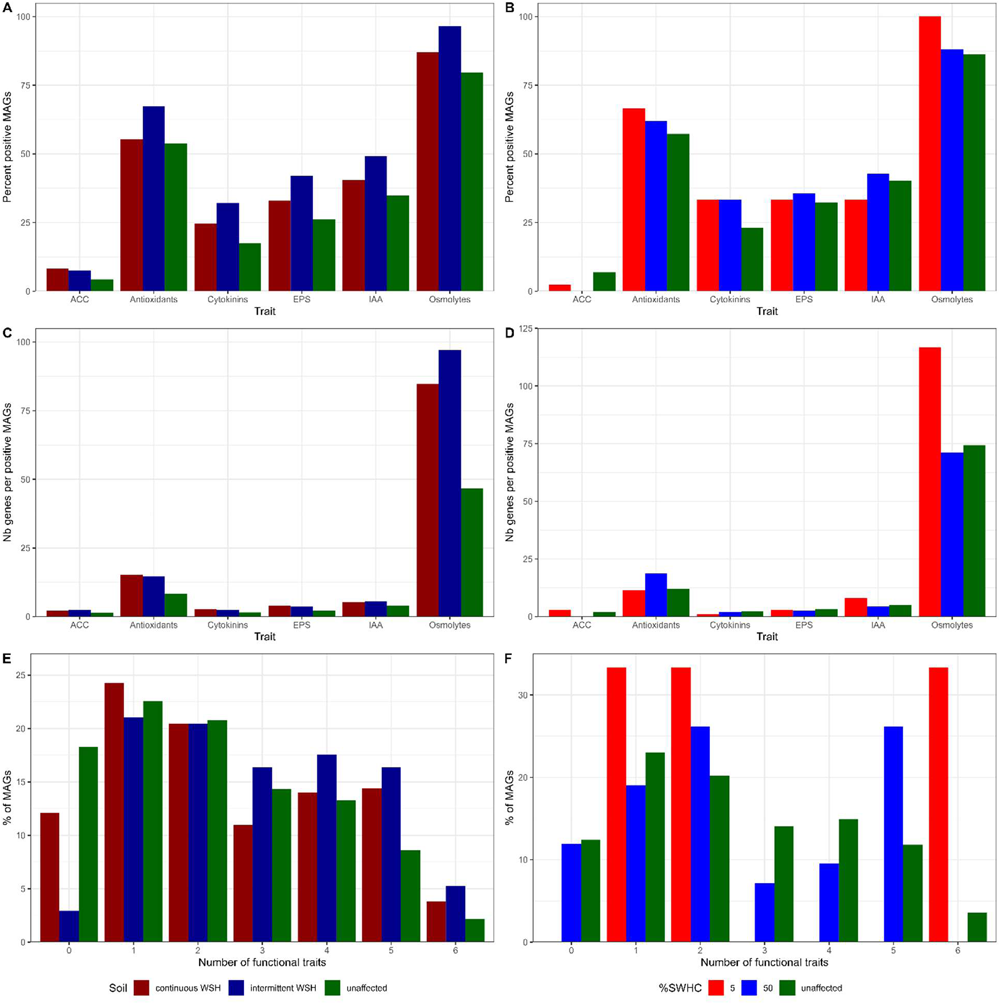
Functional traits in metagenome assembled genomes (MAGs). A and B) Percentage of the significantly affected MAGs harboring at least one copy of the genes encoding for drought-related functional traits, C and D) number of genes encoding for a drought-related functional trait per MAG, for MAGs harboring at least one of such gene, E and F) number of different traits harbored by the significantly affected MAGs. MAGs were assembled from rhizosphere samples taken from wheat growing in soil with an intermittent or continuous water stress history and subjected to low (5% soil water holding capacity) or high (50% soil water holding capacity) water availability. A, C, D: MAGs that were significantly affected by soil water stress history; B, D, F: MAGs that were significantly affected by soil water content. Since the values here are a list of MAGs that resulted from an ANOVA, statistical significance cannot be tested for. The green bar represents the unaffected MAGs for the treatment under consideration.

For the 45 MAGs affected by the actual soil water content, 3 were relatively more abundant in the LWC pots and 42 were relatively more abundant in HWC pots. The three MAGs enriched in the LWC pots all harbored genes related to osmolyte production (Fig. 3b), and each MAG had over 100 copies of them, on average (Fig. 3d), suggesting the crucial nature of this functional trait for life under low soil water content. Among these three MAGs, one had all six functional traits assessed whereas the two others had either one or two traits (Fig. 3f). The MAGs that were more abundant in HWC pots slightly more often harbored functional genes of interest than the unaffected MAGs (Fig. 3b). Except for the antioxidant genes, these MAGs, however, had less copies of the functional genes than the unaffected MAGs (Fig. 3d). As for the number of different functional traits found in the MAGs, there was not a clear trend (Fig. 3f).

## Discussion

We compared the response of wheat and its microbiota to water stress when growing in soil with almost a 40-year history of contrasting water stress frequency. In line with our hypothesis, we showed that the soil subjected to intermittent water stress better mitigated wheat fresh biomass loss in response to reduced soil water content, probably because it was enriched with microorganisms with traits beneficial for plants under water stress. These microorganisms were not only more prevalent in the soil with an intermittent history of water stress, but also were more likely to combine several beneficial traits and to have more copies of the genes encoding for these traits.

Previous exposure to stress was shown to generate a microbiota that is better adapted when facing this stress again [21, 22], and this extends to beneficial services to plants. For instance, trees grown with a microbiota with a history of stress do better when facing the same stress [23], and *Brassica rapa* better resists water stress when grown in soil that was pre-exposed to water stress [24]. Here, we showed that the frequency of stress is also important. The soil microbiota with an intermittent exposure to water stress better mitigated wheat biomass loss under low water content than the microbiota constantly exposed to water stress. Models showed that constant stressful conditions select for a microbial community dominated by a few specialists, increasing its sensitivity to environmental change and reducing its functional performance [12]. Intermittent water stress, in contrast, selected for microbial taxa that could grow at low and high water availability, i.e., generalists [12]. Experimentally subjecting a sulfidic stream microbiome to oxic/anoxic changes similarly selected for generalists active under both conditions [13]. Our data agrees well with these modelling and experimental data: MAGs enriched in the intermittent water stress history soil combined more frequently multiple functional genes related to drought resistance traits, suggesting they are generalists.

Harboring multiple traits is crucial for microorganisms to help plants during water stress. For instance, if a microorganism can mitigate plant response to stress, e.g., through interference with plant hormones, but cannot itself adapt to low soil water content, e.g., through the proficient production of osmolytes, then it will not be able to help plants during drought. Among the MAGs that were more abundant in the soil with an intermittent waster stress history, 49.1% (84/171) harbored both osmolyte and IAA genes, as compared to 40.5% (107/264) and 34.8% (97/279) for the continuous water stress history and unaffected MAGs, respectively. Microbes combining these two traits could resist low water content and, at the same time, promote plant growth. IAA and osmolyte production were the traits most influenced by water stress history frequency, being more abundant, prevalent, and diversified in the soil with an intermittent water stress history.

Osmolyte production was the most widely distributed trait in the microbial community, with 80-95% of the MAGs harboring genes of this category. Osmolyte production is one of the major mechanisms that microorganisms use to resist drought – it was estimated that 3-6% of the total annual net primary production of a grassland ecosystem is used for that purpose by microbes during a drought event [25].

We previously identified bacterial and fungal osmolyte-related genes among the most differentially expressed genes in the wheat rhizosphere under reduced soil water content [19]. Here, osmolyte production was the trait that showed the largest response to water stress history, being more prevalent, redundant, and abundant in the intermittent water stress history soils. As bacteria can transfer osmolytes to plants [5, 6], we would have expected the intermittent water stress history soil to result in a better water retention in plant tissues. This seems, however, not to have been the case. Under reduced water availability, plants growing in the intermittent water stress history soil did lose less aboveground biomass than in the continuous water stress history soil, but this was not mirrored in the leaf water content, which was unaffected. Mechanisms related to water retention, such as osmolyte and EPS production, might therefore not have played a central role in mitigating wheat biomass losses in the intermittent WSH soil. As mentioned above, osmolyte production alone might not be sufficient to mitigate water stress in plants, and the combination with other traits could be required.

The functional traits of interest were often represented by several different genes in the MAGs, especially in the ones that were more abundant in the intermittent water stress history soils. In some cases, it was because the trait in question could be encoded by various pathways or genes. One such example that was discussed above is the osmolyte production trait, which is encoded by many genes/pathways. In other cases, such as the ACC deaminase gene, redundancy indicates that the bacteria harbored several versions of the same genes. One way that bacteria can adapt to changing environmental conditions is by having two or more copies of a gene, dubbed ecoparalogs, each encoding for a protein having a different environmental optimum [26]. The accumulation of multiple ecoparalogs would be advantageous under intermittent water stress. For instance, the ACC deaminase gene was represented by an average of 2.54, 2.27 and 1.42 copies per MAG for the intermittent, continuous and unaffected MAGs, respectively. Exposure of soils to intermittent water stress seems to have selected MAGs that were containing more paralogs of genes involved in water stress response.

Although soil water content dictated the plant’s fresh biomass and leaf water content, it did not affect the microbial community structure. In contrast to the large influence of water stress history, actual soil water content only influenced the relative abundance of a few MAGs. Water stress normally decreases soil respiration [16] and microbial richness [27], increases the fungal:bacterial ratio [28], and shifts the microbial transcriptome [19]. We had reported for these soils that water stress history constrained the response of microbial communities to actual water stress [14–16]. Alternatively, the timeframe of our experiment might have been too short for these changes to translate to shifts in functional and taxonomical composition of the microbial community. In all cases, this could lead to an uncoupling of the plant-microbe interactions as the two partners do not share the same environmental cues for their response to short-term stress.

Overall, we showed that a 40-year history of intermittent soil water stress selects microbial generalists that combine important traits for plant and microbial adaptation to low soil water availability. These generalists better mitigated the effects of water stress on wheat, with plants growing in their presence having higher fresh biomass under low soil water content. Microbial generalists – because they can perform many functions in a wide range of environmental conditions – could be a key group of microorganisms for microbially-mediated plant stress resistance. We now have a clearer target for our microbial community manipulation efforts, toward improving crops’ resistance to environmental stresses.

## Acknowlegdments

We would like to thank Karelle Rheault, Éloïse Adam-Granger and Deanna Chinnerman for helping in setting up, maintaining, and sampling the experiment.

## Notes

### Competing Interest Statement

The authors have declared no competing interest.

